# Goat land degradation reduces richness and reshapes the functional composition of an ant community in a tropical dry forest

**DOI:** 10.64898/2026.03.18.712513

**Authors:** Pamela Gusmán-Montalván, Diego P. Vélez-Mora, Pablo Ramón, Elizabeth Gusmán-Montalván, Diego Dominguez, David A. Donoso

## Abstract

Tropical dry forests are among the world’s most threatened ecosystems, having lost over 60% of their original extent due to agricultural expansion and livestock grazing. Despite a growing body of research on the effects of deforestation and habitat fragmentation, the consequences of overgrazing on ant communities remain poorly documented, particularly in the Tumbesian bioregion, where uncontrolled goat grazing continues to be the primary disturbance with detrimental effects to plant communities and all the life they support.

We sampled ant communities across a goat grazing gradient in a Tumbesian dry forest in Loja, Ecuador. Nine sampling clusters were distributed across a 20 × 25 km landscape on three levels of grazing degradation (low, moderate, and high). Within each cluster, we established three 60 × 60 m plots separated by at least 250 m from each other. Eighteen pitfall traps were deployed per plot. Ant community responses to goat grazing were assessed using abundance- and incidence-based analyses and quantified vegetation structure with satellite-derived NDVI.

We collected 47,473 individuals (abundance) in 1,871 collection events (incidence) representing 22 species in six ant subfamilies. Species richness declined with grazing degradation (low = 20 spp.; moderate = 15; high = 13), and generalist forager species dominated the more degraded landscape. Beta diversity decomposition revealed a shift from turnover-dominated dissimilarity at the low moderate degradation transition to nestedness-dominated dissimilarity at the moderate high transition, indicating increasing habitat simplification with higher land degradation.

While changes in community composition showed a non-significant trend along the land degradation gradient, our trait-based analysis evidenced effects on the functional space occupied by ants: under High degradation level intensity, ants became smaller, smoother and darker.

These findings underscore the conservation value of pristine and low degradation forests and support grazing exclusion as a precautionary component of restoration strategies.

## INTRODUCTION

Tropical dry forests (TDFs) are important ecosystems that sustain half of the human population (Murphy & Lugo 1986). TDFs are among the ecosystems most affected by anthropogenic disturbance (Janzen 1988; Miles et al. 2006;), having lost more than 60% of their original extent through conversion to agricultural land and livestock grazing (Tapia-Armijos et al. 2015). Along a gradient of cumulative goat grazing, forests lose understory vegetation, soils become compact, and the forest canopy opens (Carbajal-Morón et al. 2017; Brito-Almeida et al. 2025; Temoche et al. 2025). In particular, the Tumbesian TDF of southwestern Ecuador and northwestern Peru, free-ranging goats are a dominant grazing agent, where their browsing behavior and tolerance of arid conditions make them a detrimental agent for these ecosystems (Cueva-Ortiz et al. 2019; Patiño et al. 2021). But how the impact of goat grazing on ecosystems translates to insects’ communities is subject to investigation.

Ants are one of the most abundant and ubiquitous organisms on TDFs and provide us with ecosystem services such as predation, seed dispersal, and nutrient cycling (Domínguez et al. 2016; Lattke et al. 2016; Tiede et al. 2017). Ant communities are structured by the availability of nesting sites, shade, and leaf litter, resources that decline progressively with increasing grazing (Ríos-Casanova et al. 2006; Donoso et al. 2010; Achury et al. 2012; Metz et al. 2026). These characteristics make ants particularly useful as bioindicators of habitat quality and climate change, reflecting changes in vegetation composition, functional structure, and degree of anthropogenic disturbance across a wide range of biomes (Tiede et al. 2017; Hoenle et al. 2022; Hoenle et al. 2023) and time scales (Donoso 2017; Donoso et al. 2022; Uquillas et al. 2025). Furthermore, ant morphology is tightly linked to physiological demands, i.e., thermal tolerance, desiccation resistance, and microhabitat use (Clusella-Trullas et al., 2007; Bujan et al. 2016; Law et al., 2020; Roeder et al. 2021), making trait-based approaches particularly informative for detecting ant responses to grazing via a restructuring of plant communities (Oliver et al. 2016; Eldridge et al. 2020).

However, ant responses to grazing disturbance in the Tumbesian bioregion have received limited attention, with local studies focusing primarily on the characterization of ant faunas (Domínguez et al. 2016; Lattke et al. 2016). To move forward, here we present an analysis of ant community responses to a gradient of goat-driven land degradation in the dry forest of Zapotillo, Ecuador. We tested three hypotheses: (H1) increasing land degradation reduces ant species richness, reflecting a process in which grazing filters out rare, habitat-specialist taxa; (H2) community composition shifts along the gradient through a mechanistic transition from species replacement at moderate land degradation to deterministic species loss at high land degradation; and (H3) land degradation reduces the function of ant communities. Tumbesian TDFs combine exceptional plant endemism with accelerating deforestation (Best & Kessler 1995; Aguirre & Kvist 2005), offering a well-suited system for examining how goat grazing disturbance reshapes biological communities.

## MATERIALS AND METHODS

### Study area

The study was conducted in the Tumbesian TDF of Zapotillo canton, Loja Province, southern Ecuador (4.338°S, −80.173°W; 200 to 600 m elevation; Figure 1). The climate is semiarid with a mean annual temperature of 26 °C and a mean annual precipitation of 361.7 mm, distributed bimodally with a dry (May–November) and wet (December–April) season (Espinosa et al., 2018). The woody community is dominated by drought-deciduous species characteristic of the Tumbesian floristic province, including *Bursera* grav*eolens*, *Loxopterygium huasango*, and *Ceiba trichistandra*.

**FIGURE 1.** Spatial distribution of the nine sampling clusters across the study area, Zapotillo, Loja Province, southern Ecuador. Symbols indicate land degradation: Low degradation; (Clusters 8, 10, 11), Moderate degradation; (Clusters 4, 15, 16), High degradation; (Clusters 6, 7, 9). Within each cluster, three 60 × 60 m plots were established in an L-shaped configuration (250 m edge-to-edge separation). The three disturbance types are spatially intermingled across the landscape. Scale bar = 6 km. Inset shows the location of the study area within Ecuador. Inset photographs illustrate representative vegetation structure at each land degradation level.

Land use in the region has undergone substantial change over the past three decades (Patiño et al. 2021). Selective logging of valuable timber species reduced canopy cover across much of the coastal dry forest until protective legislation was enacted in 1978, after which timber extraction declined but did not cease entirely (Tapia-Armijos et al., 2015). Between 1978 and 2008, the broader region lost approximately 33% of seasonally dry forest and 18% of scrubland cover, predominantly converted to pastureland (Tapia-Armijos et al., 2015). More recently, the construction of the Zapotillo irrigation system in 2011 expanded agricultural land and increased the number of free-ranging goat and cattle herds grazing in adjacent forest remnants.

### Sampling design

In 2019, we established nine sampling clusters across a landscape spanning approximately 20 × 25 km, representing three levels of cumulative grazing over decades based on vegetation cover and canopy structure: low (minimal- or no grazing; intact canopy; Clusters 8, 10, 11), moderate (Clusters 4, 15, 16), and high (decades of sustained, intensive grazing; sparse canopy and degraded understory; Clusters 6, 7, 9) (Figure 1 and Table S1). Plot setting and categorization, and cluster terminology, were previously reported in a larger study that focused on plants responses to goat grazing (see Cueva-Ortiz et al. 2019). The three levels of land degradation were spatially intermingled. Clusters of different land degradation occur as close as 1.86 km apart (e.g., Clusters 11–15), while clusters of the same land degradation can be up to 19.42 km apart (e.g., Clusters 10–11). Level centroids were separated by 5.21–8.93 km, well within the geographic range occupied by any single level (Figure 1; Table S1). Within each cluster, we arranged three 60 × 60 m plots in an L-shaped configuration with 250 m edge-to-edge separation between adjacent plots, yielding a total of 27 plots. In this two-level nested structure (plots within clusters, clusters within land degradation levels), the cluster constitutes the unit of treatment replication (n = 3 clusters per degradation level), while the plot is the unit of ecological observation (9 clusters × 3 plots = 27 plots). We deployed 486 pitfall traps in total (3 degradation levels × 3 clusters × 3 plots × 18 traps = 486 traps).

To evaluate whether the spatial arrangement of sampling plots could confound our results, we applied two complementary approaches. First, partial Mantel tests (Mantel, 1967; vegan; Oksanen et al., 2022; 999 permutations) compared pairwise geographic distances with Bray-Curtis (raw ant abundance) and Jaccard (ant incidence) compositional dissimilarity matrices, controlling for degradation level as an ordered matrix at both the plot level (n = 27; 351 pairs) and the cluster level (the true unit of treatment replication; n = 9; 36 pairs). Second, spline correlograms (ncf; Bjørnstad, 2020; 999 bootstrap permutations) were computed on the first residual axis of a distance-based redundancy analysis (dbRDA; Bray-Curtis; vegan), representing compositional variation not explained by the gradient.

### Vegetation gradient

For our analyses, we also used vegetation status along the gradient as a predictor, using the Normalized Difference Vegetation Index (NDVI; Tucker, 1979) for each of the 27 plots from Sentinel-2 satellite imagery (acquisition date: November 2018, 10 m resolution) processed with Google Earth Engine (Gorelick et al., 2017) via the rgee v1.1.8 package (Aybar et al., 2020). Plot centroids, recorded in UTM Zone 17S (EPSG:32717), were reprojected to geographic coordinates (EPSG:4326) using the sf package (Pebesma, 2018). A circular buffer of 10 m radius was defined around each centroid. We used the COPERNICUS/S2_HARMONIZED collection (Level 1C) filtered to November 2018, coinciding with the field campaign, and excluded images with cloud cover exceeding 20%. NDVI was computed as (ρB8 − ρB4) / (ρB8 + ρB4), where ρB8 is near-infrared (842 nm) and ρB4 is red (665 nm). Mean NDVI per plot was obtained from 2–3 valid images.

### Current goat grazing

Goat herding is the dominant land use in the study area and creates the gradient of cumulative grazing examined in this study (Cueva-Ortiz et al. 2019; Patiño et al. 2021). As a measure of current total livestock activity, we used manure deposition (dry weight) along a 1 × 60 m transect located at the edge of each 60 × 60 m plot. Besides goat manure, this measure includes also cow and horse manure which was also present throughout the study area but in lower amounts.

### Ant species identification

All specimens were identified to species (or morphospecies). Due to the lack of identification literature on Ecuadorian ants (Salazar et al. 2015; Dominguez et al. 2016), we mainly relied on keys for Costa Rican and Colombian ants (e.g., Fernández et al., 2019). Voucher samples are deposited in the MUTPL museum collection at the Universidad Técnica Particular de Loja. For genera with difficult taxonomy (like *Solenopsis* and *Nylanderia*) we preferred to be conservative and merge all individuals and species under the genus umbrella. All species were delimited and coded consistently by the same observer throughout the study to minimize intra-study inconsistency.

### Functional traits

Six functional traits were measured for all ant species recorded in this study related to ant foraging strategies and life history (Silva 2010, Arnan et al. 2014, Yates et al. 2014, Gibb et al. 2015, Bishop et al. 2016, Ohkawara et al. 2017). Traits chosen for our analysis were based on the framework by Parr et al. (2017) and include head length (HL), used as a proxy for body size and for foraging and resource-handling capacity, cuticular sculpturing (SCUL), scored on an ordinal scale from 1 to 3 (smooth to strongly sculptured surface), and pilosity (PIL), quantified as the number of standing hairs present on the pronotum; both traits are associated with desiccation resistance and physical protection against adverse microclimatic conditions. Finally, ant coloration was characterized based on the dominant color of the head (DCH), mesosoma (DCM), and gaster (DCG), coded following a standardized color scale (1–24). For each species, at least 5 individuals were measured (45x magnification, with a measurement error of 0.01 mm) using an OLYMPUS SZ6 stereomicroscope (Table S5).

### Statistical Analyses

#### Vegetation gradient

NDVI differences among degradation levels were tested with a linear mixed model (lmer: NDVI_mean ~ degradation + (1|Cluster)), with Cluster as a random intercept to account for the nested plot structure; post-hoc pairwise comparisons used emmeans with Satterthwaite degrees of freedom and Holm correction.

#### Ant abundance and incidence data

In this study we used both abundance and incidence data. Abundance corresponds to the total number of individuals by species trapped by our design, while incidence corresponds to the presence/absence of species in the traps and is more indicative of colony number. Contrasting abundance with incidence data allowed us to assess whether community patterns were robust to the numerical dominance of a few abundant species. However, because pitfall traps capture foraging individuals over a defined period, abundance data from this method reflect activity density rather than a direct census of colony or worker density (Andersen, 1991; Parr & Chown, 2001), and raw abundance values reported in this study should therefore be interpreted with this caveat in mind; we place greater analytical weight on incidence-based results where the two approaches diverge.

#### Sample completeness and ant alpha diversity

Ant alpha diversity at the plot level (n = 27) was quantified using Hill numbers for orders q = 0 (species richness), q = 1 (Shannon diversity), and q = 2 (Simpson diversity). Sample completeness was evaluated using rarefaction and extrapolation curves (q = 0) implemented in iNEXT v. 3.0 (Hsieh et al., 2016), with pooled abundance vectors per level as input. Asymptotic richness was estimated with Chao1 and ACE (abundance; Chao et al., 2014; vegan; Oksanen et al., 2022) and Chao2 (incidence). Differences among degradation levels in Hill numbers were tested with linear mixed models (lmer: qD ~ degradation + (1|Cluster), lme4/lmerTest; Satterthwaite df), with Cluster as a random intercept to account for the nested plot structure. Post-hoc pairwise comparisons used emmeans with Holm correction.

#### Community composition

We visualized ant community composition using an NMDS ordination computed on all 27 plots. However, because the cluster is the unit of treatment replication, statistical tests were conducted at the cluster level (n = 9; 3 per level). For abundance data, we used the Hellinger-transformed Bray-Curtis dissimilarity matrix. For incidence data, we used the Jaccard dissimilarity matrix. To visualize the NDVI gradient in ordination space, we projected the NDVI vector onto the NMDS using envfit (999 permutations) computed on the nine cluster centroids in NMDS space (mean NMDS coordinates of the three plots per cluster), ensuring that the permutation test was conducted on independent replication units. We tested compositional differences among degradation levels using PERMANOVA (adonis2, 999 permutations) on cluster matrices: mean abundances per cluster (Hellinger-transformed) for abundance data, and binary presence– absence matrices (species recorded in ≥ 1 plot per cluster) for incidence data. For PERMANOVA and SIMPER we used Sørensen dissimilarity for incidence data and Bray-Curtis for abundance data, ensuring full consistency across multivariate tests. Multivariate dispersion was assessed with PERMDISP (betadisper/permutest, 999 permutations) on the same cluster matrices using square-root-transformed dissimilarities, which are Euclidean and avoid negative eigenvalues in the underlying principal coordinates analysis (Anderson et al. 2006). Owing to the inherent collinearity between land degradation and NDVI— which both proxy the same vegetation dynamics—we omitted a joint PERMANOVA model. Instead, we used variation partitioning (varpart, vegan) to quantify the variance in community composition uniquely explained by land degradation, uniquely explained by NDVI, and shared between the two predictors. The significance of unique fractions was tested using conditioned RDA (rda) for abundance data and conditioned db-RDA (capscale) for incidence data (999 permutations each). SIMPER analysis identified species contributing most to pairwise dissimilarity among land degradation levels (cumulative contribution ≤ 70%), computed on cluster matrices.

Beta diversity was decomposed into spatial turnover (species replacement, β.SIM) and nestedness-resultant dissimilarity (species loss, β.SNE) following Baselga (2010), using the betapart package (Baselga & Orme, 2012). A cluster-level presence-absence matrix was used as input (a species scored as present if detected in ≥ 1 plot within the cluster). Both global multi-site and pairwise decompositions were computed using the Sørensen family index, and mean component values were calculated for all cluster pairs belonging to different land degradation levels

#### Functional trait analysis

Community-weighted means (CWM) of each trait were calculated as abundance-weighted averages across all genera present in each plot. Functional diversity was quantified using the dbFD function in the R package FD (Laliberté & Legendre, 2010), which computes Gower distances among species based on mixed trait types and returns four complementary indices: functional richness (FRic), functional evenness (FEve), functional divergence (FDiv), and functional dispersion (FDis). Negative eigenvalues in the underlying PCoA were corrected using the Cailliez method.

To test the effect of land degradation level on functional diversity indices and CWM traits, we fitted linear mixed models (LMMs) with degradation as a fixed factor and cluster as a random intercept, using the lmerTest package (Kuznetsova et al., 2017). Significance was assessed via Type III F-tests with Satterthwaite’s degrees of freedom. For continuous predictors (total livestock manure, tree species diversity, Simpson; normalized difference vegetation index, NDVI), we compared models using AICc (Burnham & Anderson, 2002).

#### Multivariate functional composition

To explore how land degradation and vegetation structure jointly shape community functional composition, we performed a principal component analysis (PCA) on the standardized CWM values of the six selected traits, aggregated to the cluster level (n = 9, the true unit of replication). We then projected three environmental predictors onto the PCA ordination space using the envfit function (vegan package; Oksanen et al., 2022) with 999 permutations: total livestock manure, normalized difference vegetation index (NDVI), and tree species diversity (Simpson index). Because envfit tests the squared correlation coefficient (r^2^), which requires greater sample sizes to detect significance than simple correlations, we also report Pearson correlation coefficients between each predictor and the first two principal components, with associated p-values.

To visualize the species-based functional space and its contraction along the grazing gradient, we computed Gower distances on the same six traits and performed a principal coordinates analysis (PCoA) with Cailliez correction for negative eigenvalues, using the pcoa function in the ape package (Paradis & Schliep, 2019). Species positions in the PCoA ordination were displayed with bubble sizes proportional to their total abundance within each land degradation level, allowing visual comparison of the occupied functional space across treatments. All analyses were conducted in R v. 4.3.1 (R Core Team, 2021).

## RESULTS

### Spatial independence of the survey plots

Partial Mantel tests indicated that geographic proximity of plot (Bray-Curtis: r = 0.055, p = 0.238; Jaccard: r = 0.059, p = 0.211) and cluster (Bray-Curtis: r = −0.071, p = 0.589; Jaccard: r = −0.029, p = 0.548) did not predict compositional similarity once land degradation level is accounted for. Spline correlograms showed positive autocorrelation only at distances below 0.5 km (corresponding to within-cluster plot spacing), with the 95% bootstrap confidence band including zero at all inter-cluster distances (> 0.5 km; Figure S1).

### Vegetation gradient

NDVI differed significantly with land degradation (F□,□ = 8.31, p = 0.019). High plots had lower NDVI than both Low (p = 0.031) and Moderate plots (p = 0.031), while Low and Moderate plots did not differ (p = 0.755; Table S2) among themselves. Cluster means ranged from 0.083 (Cluster 6, High) to 0.196 (Cluster 15, Moderate; Figure S2).

### Ant species

A total of 47,473 individuals in 1,871 collection events were collected representing 22 species from 6 subfamilies. The total number of individuals varied with land degradation (Low: 17,542; Moderate: 10,628; High: 19,289). Species richness declined across the gradient, from 20 species in Low degradation level, to 15 in Moderate, to 13 in High.

Low degradation plots harbored several species not recorded elsewhere (*Carebara* sp., *Cephalotes maculatus, Leptogenys* sp1 and *Platythyrea* sp1) (Figure 2a, 3b). Moderate degradation had no exclusive species, while High degradation level yielded one unique species: *Cardiocondyla* sp1. In contrast, the incidence of *Crematogaster* spp. increased from 77.8% under Low to 100% under High degradation level (Figure 2b).

**FIGURE 2.**
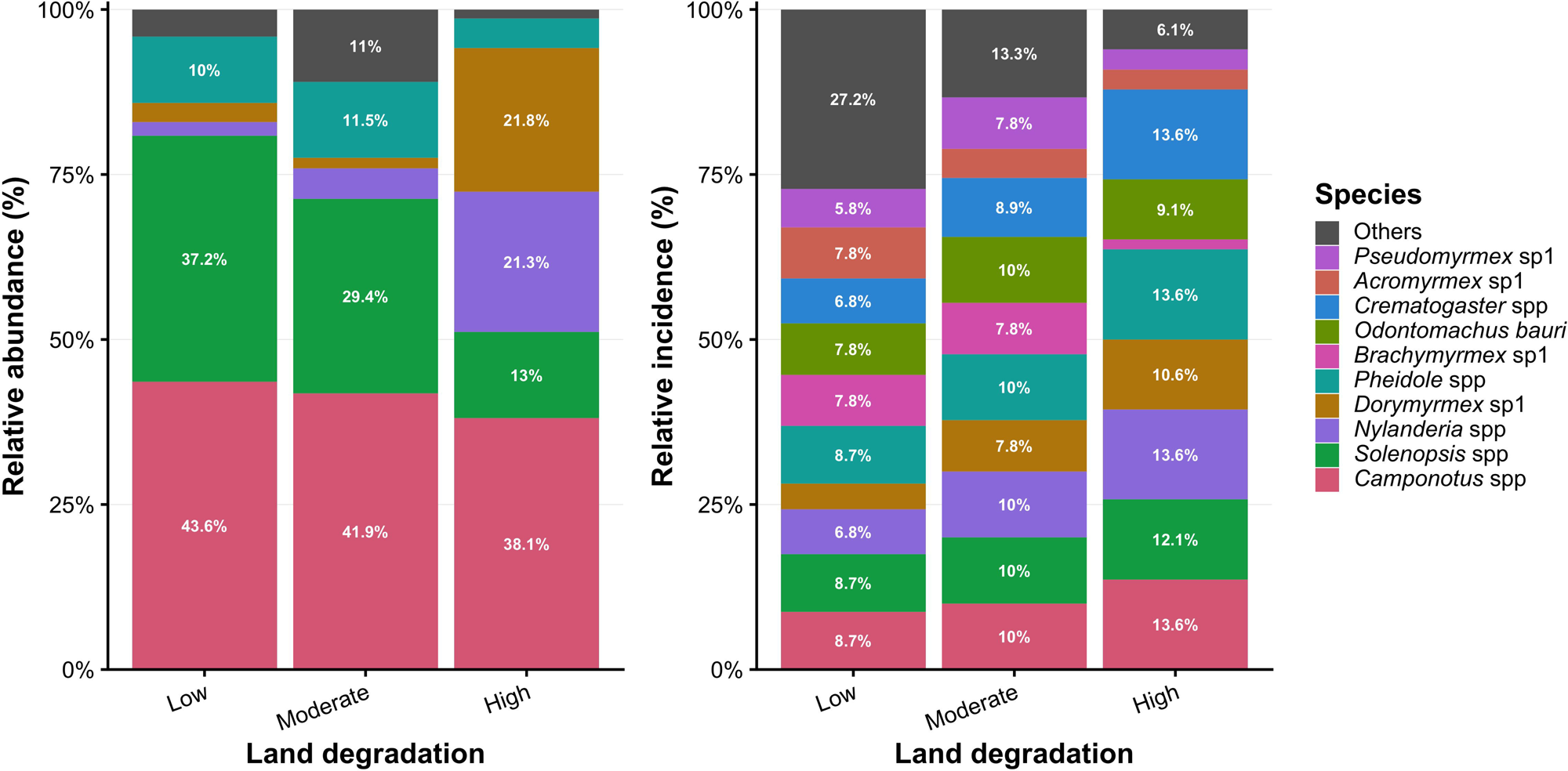
Ant community composition across the grazing gradient in the Tumbesian TDF of Zapotillo, Ecuador. (a) Relative abundance (% of total individuals) per species. (b) Relative detection frequency (% of total plot-species detection events). Land degradation levels: Low degradation, Moderate degradation, High degradation.

### H1. Alpha diversity and sampling completeness

Rarefaction curves approached the asymptote for all land degradation levels in both abundance- and incidence-based analyses (Figure 3a, 3c), and asymptotic estimators closely matched observed richness (Low degradation level: S.obs = 20; Moderate: S.obs = 15; High: S.obs = 13; Figure 3b), indicating adequate sampling completeness. Incidence-based Chao2 estimates (20, 15.9 and 13 for Low, Moderate and High degradation Levels, respectively; Figure 3d) confirmed this pattern independently of raw individual counts.

**FIGURE 3.**
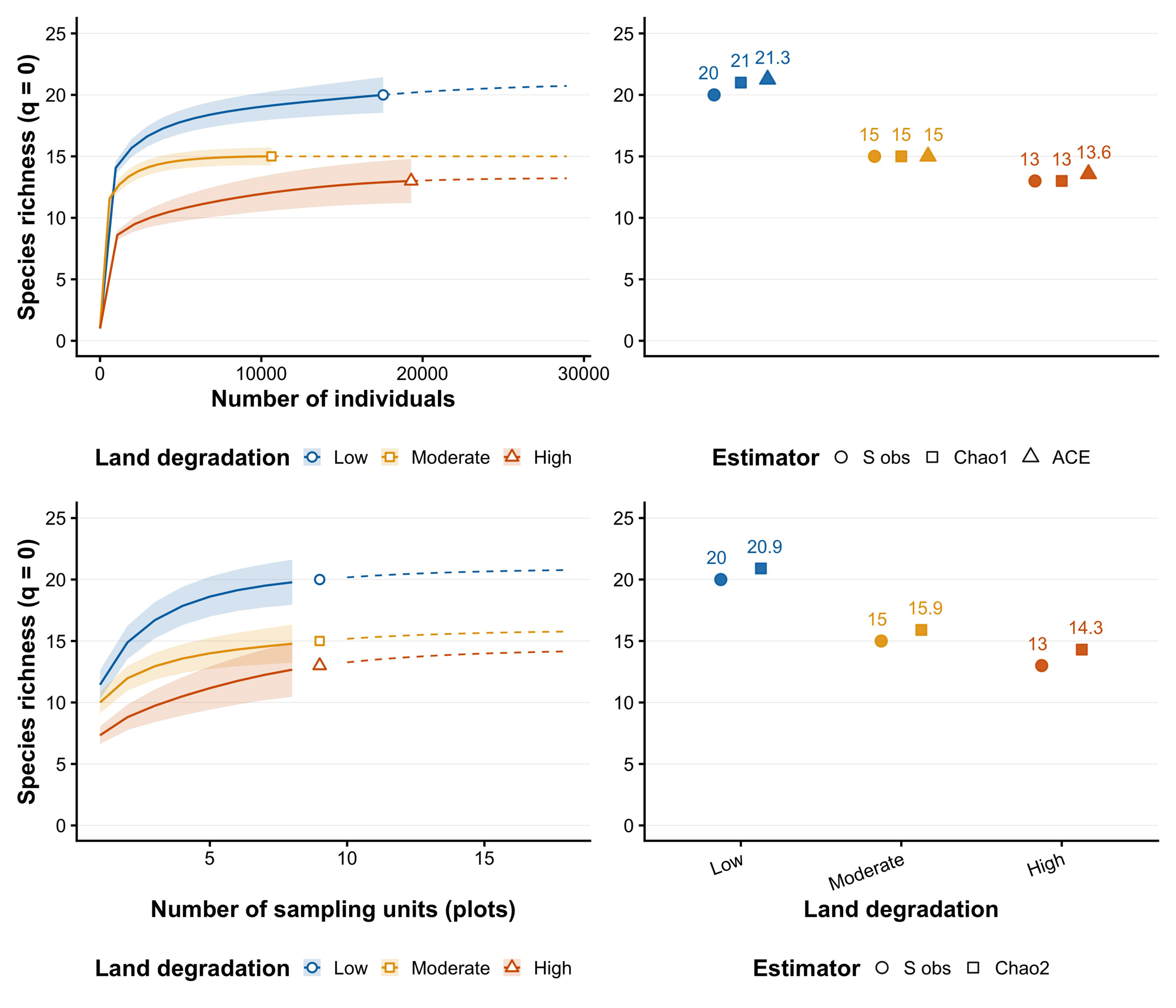
Species richness estimation across the grazing gradient in the Tumbesian TDF of Zapotillo, Ecuador. (a) Abundance-based rarefaction and extrapolation curves for species richness across land degradation levels, using individuals as the sampling unit. (b) Asymptotic richness estimators S obs, Chao1 and ACE, showing near-complete sampling. (c) Incidence-based rarefaction and extrapolation curves for species richness, using plots (n = 9 per level) as the sampling unit. (d) Observed richness (S obs) and Chao2 asymptotic estimate based on plot-level incidence data.

Linear mixed models showed a significant effect of land degradation level on species richness (q = 0: F_2,6_ = 5.64, p = 0.042), but not on Shannon (q = 1: F_2,6_ = 0.26, p = 0.779) or Simpson diversity (q = 2: F_2,6_ = 0.04, p = 0.960; Figure S3). Post-hoc comparisons did not detect significant pairwise differences after correction: Low vs. High degradation approached but did not reach significance (p = 0.052), and Low vs. Moderate degradation (p = 0.388) and Moderate vs. High degradation (p = 0.118) did not differ. These results indicate that grazing was associated with a reduction in species richness, an effect not mirrored by abundance-weighted diversity metrics.

Despite having the highest richness, Low degradation showed the lowest q = 1 diversity, while Moderate degradation showed the highest (Table S3). This pattern is consistent with strong numerical dominance by *Camponotus* spp. in more pristine forests, which reduces abundance-weighted effective diversity independently of richness.

### H2. Community composition

NMDS ordinations based on abundance (Stress = 0.122) and incidence (Stress = 0.137) data showed partial, non-significant separation of land degradation in ordination space, with High degradation clusters displaced along the NDVI gradient (Figure 7 a,b). PERMANOVA on cluster centroids (n = 9) did not detect significant compositional differences among land degradation for either abundance (F_2,6_ = 2.54, R^2^ = 0.459, p = 0.101) or incidence (F_2,6_ = 3.0, R^2^ = 0.50, p = 0.070) data. Homogeneity of multivariate dispersion (PERMDISP: F_2,6_ = 0.54, p = 0.598) or incidence (F_2,6_ = 0.45, p = 0.635) indicates that observed ordination patterns are not attributable to differences in within-group. The NDVI vector showed a positive association with ordination space for both abundance-based (r^2^ = 0.544, p = 0.096) and incidence-based (r^2^ = 0.623, p = 0.052), data though neither reached the conventional significance threshold.

Because PERMANOVA did not detect significant compositional differences, the SIMPER results below are presented as a descriptive characterization of which species contributed most to observed dissimilarity and should not be read as confirming compositional divergence among land degradation levels. Abundance-based SIMPER showed that dissimilarity was distributed across multiple generalist species, with no single taxon dominating any pairwise comparison. In the Low vs. Moderate degradation comparison, seven species together accounted for 69.4% of dissimilarity: *Solenopsis* spp (16.3%), *Camponotus* spp (15.1%) and *Brachymyrmex* sp1 (11.2%). In the Low vs. High degradation contrast, *Dorymyrmex* sp1 (20.2%), *Nylanderia* spp (18.2%), *Solenopsis* spp (14.7%) and *Camponotus* spp (10.9%) jointly accounted for 64.0% of dissimilarity. In the Moderate vs. High degradation contrast, *Dorymyrmex* sp1 (21.0%), *Solenopsis* spp (15.5%), *Nylanderia* spp (14.6%) and *Camponotus* spp (12.0%) together represented 63.1% of dissimilarity (Figure 7c).

Incidence-based SIMPER showed a different pattern, with dissimilarity distributed across species whose detection frequency declined along the grazing gradient (Figure 7d). In the Low vs. Moderate degradation contrast, six species together accounted for 65.5% of dissimilarity: *Platythyrea* sp1 (13.1%), *Cephalotes maculatus* (13.1%), *Neivamyrmex* sp1 (11.6%), *Pachycondyla harpax* (9.5%), *Azteca* sp1 (9.1%), and *Labidus* sp1 (9.1%). In the Low vs. High degradation contrast, seven species contributed nearly equally (6.8–10.3%): *Brachymyrmex* sp1 (10.3%), *Azteca* sp1 (10.3%), *Cephalotes maculatus* (9.4%), *Pachycondyla harpax* (9.4%), *Platythyrea* sp1 (9.4%), *Rogeria* sp1 (9.4%) and *Neivamyrmex* sp1 (6.8%), together accounting for 65.0% of dissimilarity. In the Moderate vs. High degradation contrast, *Rogeria* sp1 (19.9%), *Brachymyrmex* sp1 (13.7%), *Azteca* sp1 (13.4%), and *Pachycondyla harpax* (13.0%) together accounted for 60.0% of dissimilarity.

Beta diversity decomposition indicated a shift in the relative contribution of turnover and nestedness components along the grazing gradient (Figure 8, Table S4). The Low vs. Moderate degradation comparison was dominated by turnover (β.SIM = 0.099, β.SNE = 0.078, SNE/SOR = 0.41, β.SOR = 0.177). The Moderate vs. High degradation comparison showed the opposite pattern, with nestedness as the dominant component (β.SNE = 0.136, β.SIM = 0.091, SNE/SOR = 0.55, β.SOR = 0.227). The Low vs. High degradation contrast showed the highest total dissimilarity (β.SOR = 0.277), with nestedness contributing most of this dissimilarity (SNE/SOR = 0.72, β.SIM = 0.087, β.SNE = 0.190). This shift from turnover-dominated to nestedness-dominated dissimilarity along the gradient indicates that species loss, rather than replacement, increasingly accounts for compositional differences at higher grazing.

### H3. Functional diversity indices

Community-weighted means of the six selected traits showed consistent directional trends along the grazing gradient (Figure S4), several of which were statistically supported when tested against continuous grazing predictors (see below). Head length (HL) and pilosity (PIL) were lower under High rather than Low degradation. Sculpturing (SCUL) varied under Low and Moderate degradation (CWM range: 1–3) but was uniformly low (≈1.0) under High degradation. The three color-based traits (DCH, DCM, DCG) were lower under High degradation relative to Low degradation. When modelled against continuous predictors, total livestock manure was the best-supported predictor (AICc-selected) for CWM traits related to body structure: it was significantly associated with pilosity (t9.0 = −2.35, p = 0.044), sculpturing (t9.6 = −2.30, p = 0.045), and head coloration (t8.9 = −2.60, p = 0.029), and marginally associated with head length (t8.6 = −2.10, p = 0.067) and body length (t8.8 = −2.19, p = 0.057).

Functional diversity indices varied among land degradation levels (Figure 4). Functional richness (FRic) declined numerically from 0.261 ± 0.054 (mean ± SD) under Low degradation to 0.209 ± 0.066 under Moderate degradation and 0.144 ± 0.083 under High degradation, but this trend did not reach statistical significance (F_2,6_ = 4.89, p = 0.055). Functional divergence (FDiv) was similar between Low degradation (0.870 ± 0.061) and Moderate degradation (0.903 ± 0.033) and lower under High degradation (0.716 ± 0.181), also without reaching significance (F_2,6_ = 3.00, p = 0.125). Functional evenness (FEve) and functional dispersion (FDis) did not differ significantly among degradation levels (FEve: F_2,6_ = 3.12, p = 0.118; FDis: F_2,24_ = 0.93, p = 0.408). Among continuous predictors, NDVI showed a marginally significant association with FRic (t17.6 = 2.26, p = 0.037), while no predictor was significantly associated with FDis, FEve, or FDiv at the plot level (Figure S5).

**FIGURE 4.**
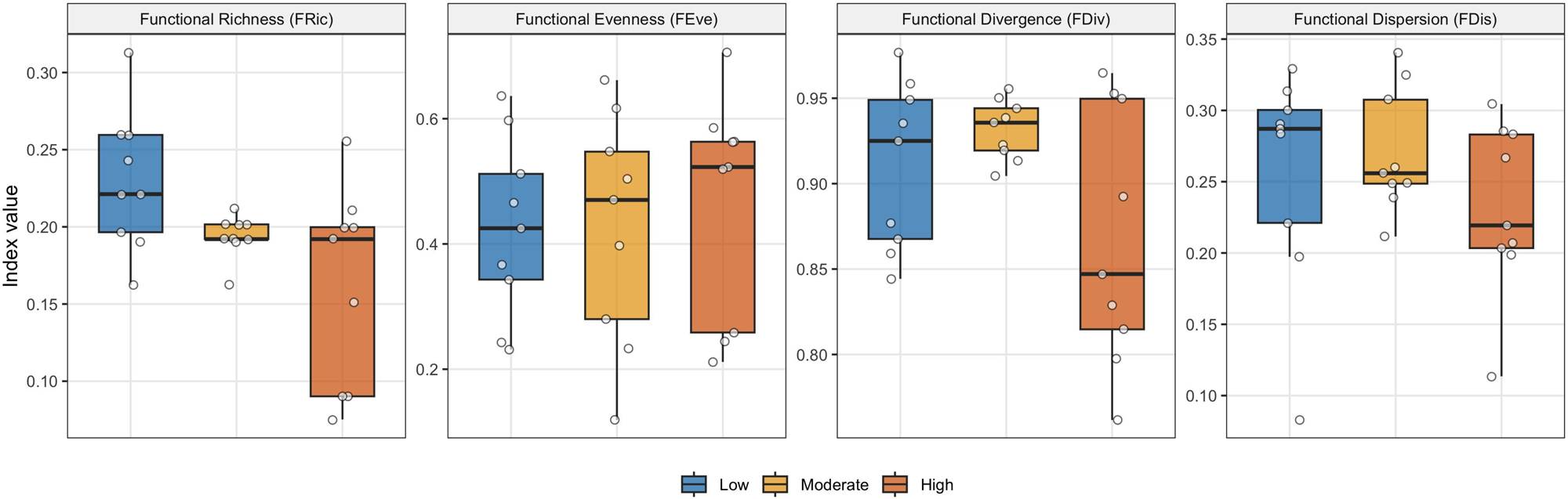
Functional diversity indices: Functional Richness (FRic), Functional Evenness (FEve), Functional Divergence (FDiv), and Functional Dispersion (FDis) across the grazing gradient. Boxplots show median and interquartile range; points show individual plot values (n = 9 per level). Land degradation levels: Low degradation, Moderate degradation, High degradation.

### Multivariate functional composition (PCA)

The PCA on cluster-level CWM of the six functional traits captured 98.4% of total variance in the first two axes (PC1: 76.6%; PC2: 21.8%; Figure 5). Projecting environmental predictors onto the PCA space identified two statistically significant relationships. Total livestock manure was significantly correlated with PC1 (Pearson r = –0.689, p = 0.040), indicating that increasing grazing is associated with communities of smaller-headed, smoother, and darker-colored genera. Tree species diversity (Simpson index) was significantly correlated with PC2 (r = 0.672, p = 0.048), indicating an association between higher tree diversity and lighter body coloration. NDVI showed a positive but non-significant association with PC2 (r = 0.594, p = 0.092), which we interpret as consistent with, but not independent confirmation of, the tree-diversity–coloration association above. Plot-level subsamples (n = 27) projected onto the cluster-level PCA space showed within-cluster variation consistent with the among-cluster gradient described above.

**FIGURE 5.**
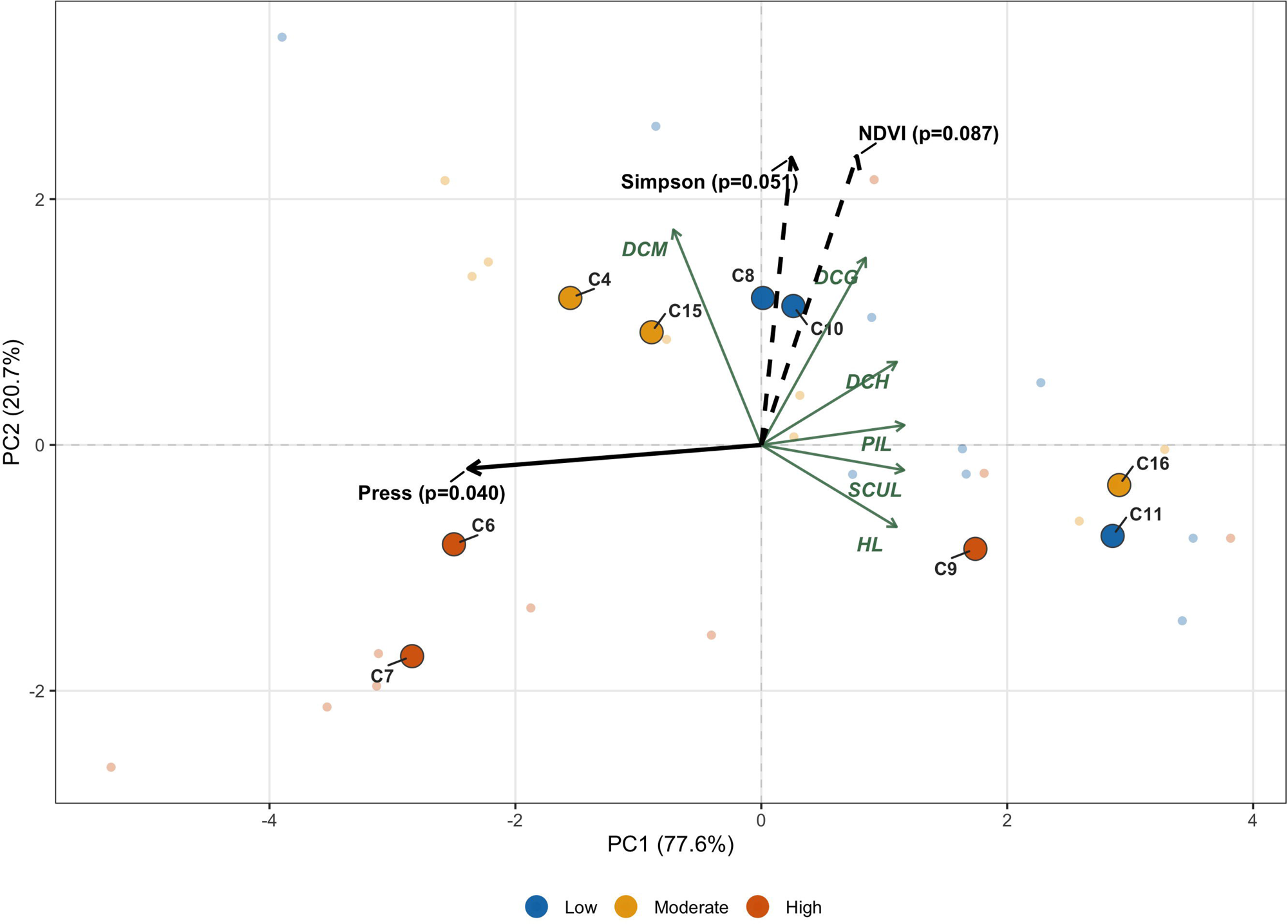
Principal component analysis (PCA) biplot of cluster-level CWM of six functional traits (n = 9 clusters). Green arrows show trait loadings; black arrows show environmental predictors projected via envfit (999 permutations). Solid arrows indicate significant associations (p < 0.05); dashed arrows indicate non-significant associations. Small translucent points show plot-level subsamples (n = 27) projected onto the cluster-level PCA space.

A supplementary PCA including all 17 traits confirmed that the six-trait subset captured the dominant functional gradients (PC1 = 84.5%; Figure S6). All six traits loaded positively on PC1 except DCM (mesosoma color; loading = –0.271), such that PC1 ordered communities from those dominated by large-headed, sculptured, pilose, light-colored genera (positive scores) to those dominated by small-headed, smooth, glabrous, dark-colored genera (negative scores). PC2 separated communities primarily by coloration, with DCM (0.699) and DCG (0.579) showing the strongest positive loadings.

### Species functional space (PCoA)

The PCoA based on Gower distances among the 22 genera captured 47.6% of total variation in the first two axes (Axis 1: 30.3%; Axis 2: 17.3%; Figure S7, and Figure 6). Axis 1 separated large-bodied, heavily sculptured genera (e.g., *Pachycondyla, Odontomachus, Cephalotes;* negative scores) from small-bodied, smooth genera (e.g., *Pheidole, Pseudomyrmex, Neivamyrmex*; positive scores). Axis 2 separated genera primarily by body coloration, with *Camponotus* (large, dark) at the negative extreme and *Dorymyrmex* and *Cardiocondyla* (light-coloured) at the positive extreme. When species occurrence was partitioned by land degradation (Figure 6), the occupied functional space became progressively smaller with increasing land degradation. All 22 species were recorded under Low degradation, including peripherally positioned taxa such as *Cephalotes, Pachycondyla,* and *Odontomachus.* Under Moderate degradation, several functionally distinct species were not recorded (e.g., *Cephalotes, Cyphomyrmex*). Under High degradation, *Pachycondyla* was not recorded, *Odontomachus* was recorded at minimal abundance, and the remaining occurrences were concentrated among the numerically dominant species *Dorymyrmex, Nylanderia* and *Solenopsis*.

**FIGURE 6.**
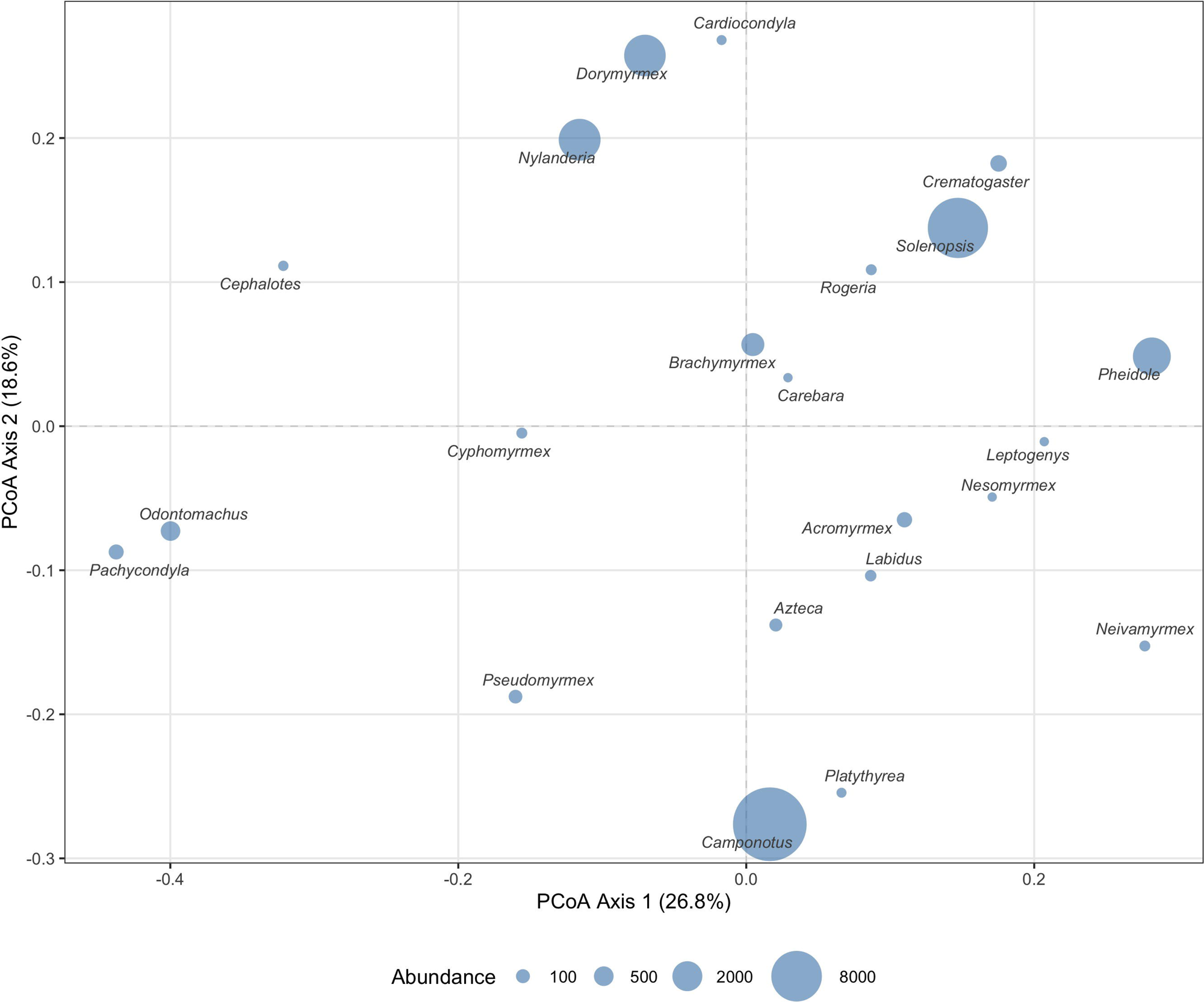
Species-based functional space based on PCoA of Gower distances among 22 genera using six functional traits, partitioned by land degradation level. Bubble sizes are proportional to species abundance within each level. The occupied functional space contracts progressively from Low degradation (all 22 morphospecie) to High degradation (dominated by four generalist genera).

**FIGURE 7.**
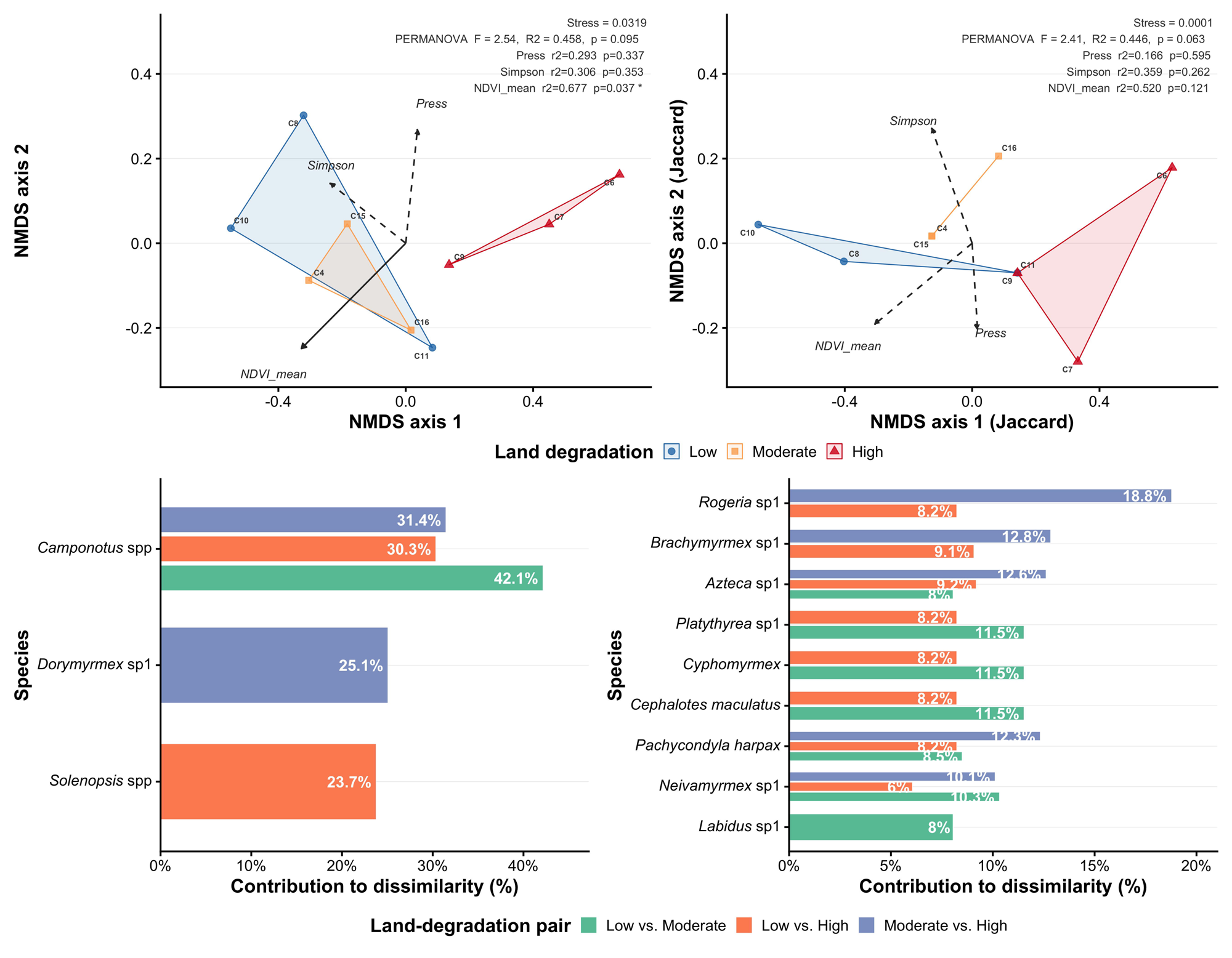
Ant community composition across the grazing gradient. (a, b) NMDS ordinations based on Bray-Curtis (abundance; stress = 0.122) and Jaccard (incidence; stress = 0.137) dissimilarities (n = 27 plots); convex hulls connect cluster centroids; arrows show the NDVI gradient (r^2^ = 0.544, p = 0.096 and r^2^ = 0.623, p = 0.052, respectively; not statistically significant). PERMANOVA on cluster centroids (n = 9) indicated did not detect significant compositional separation among land degradation levels (Bray-Curtis: F = 2.54, R^2^ = 0.459, p = 0.101; Jaccard: F = 3.00, R^2^ = 0.500, p = 0.070). (c, d) SIMPER analysis showing species contributing most to pairwise dissimilarity (up to 70% cumulative contribution) based on abundance (c) and incidence (d) data shown descriptively, not as confirmation of significant compositional divergence.

**FIGURE 8.**
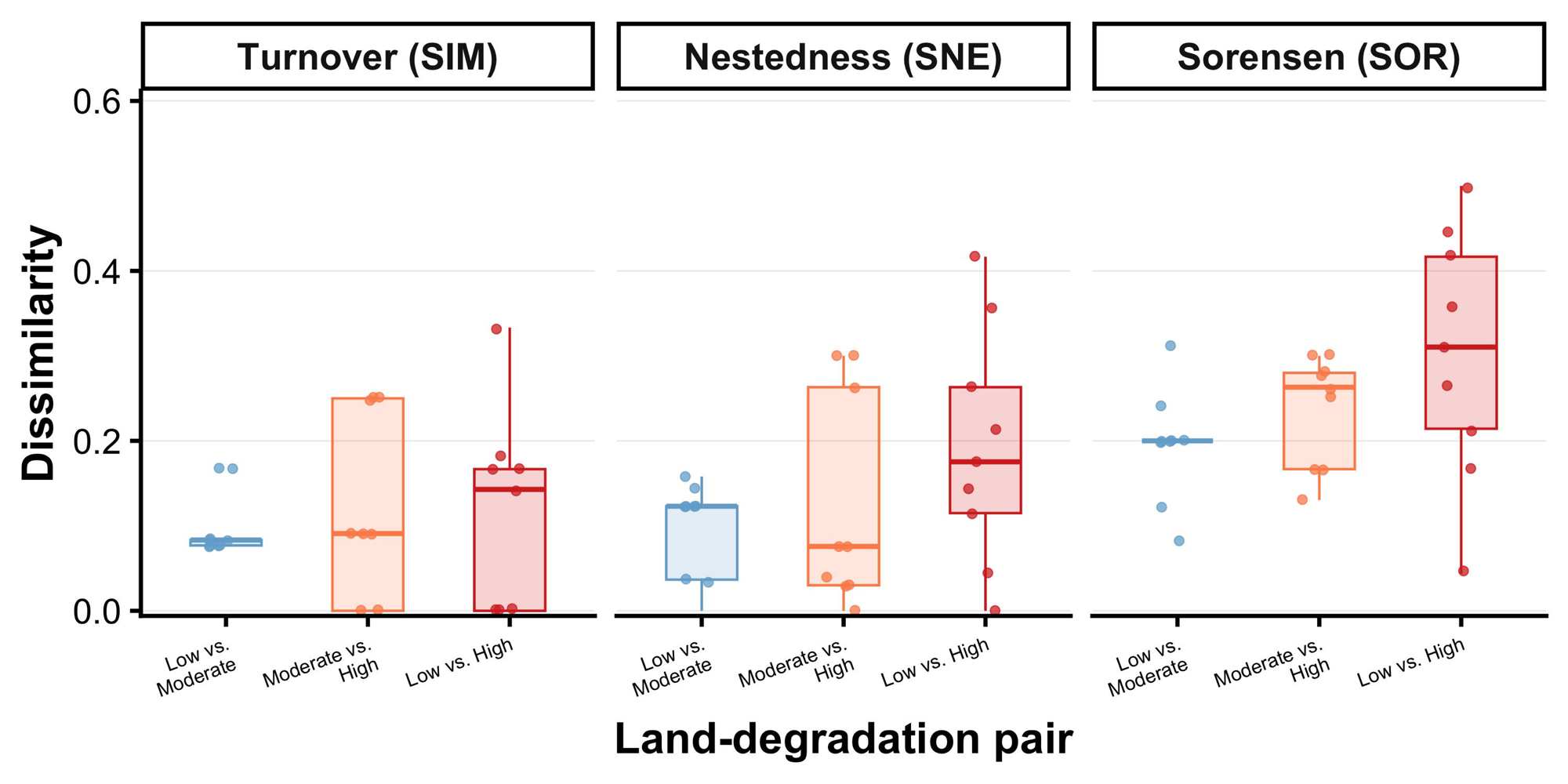
Pairwise beta diversity decomposition between land degradation levels based on cluster-level presence-absence data (n = 9 clusters, 3 per land degradation; a species scored as present if detected in ≥ 1 plot within the cluster). Each point represents one cluster pair. Mean nestedness proportions (β.SNE/β.SOR): Low vs. Moderate degradation = 0.41; Moderate vs. High = 0.55; Low vs. High = 0.72.

## DISCUSSION

In this work, we explore the responses of ant communities to increasing goat grazing intensity in the dry forest of south Ecuador. Previous work (Cueva-Ortiz et al. 2019) linked grazing intensity to more open and less-structured forests. Our results provide support for the three hypotheses tested. Grazing was associated with a significant decline in ant species richness (H1) and with a directional shift in functional trait composition supported by multiple independent lines of evidence (H3, partial support). In contrast, we found no statistically significant evidence of change in overall community composition among land degradation (H2) even though we found a shift in the process generating dissimilarity along the gradient (from turnover to nestedness).

Our results found a more diverse set of species in Low degradation plots. Here, the species we found were ecologically informative regarding the mechanisms of richness loss. *Cephalotes maculatus* is an obligate arboreal species requiring hollow twigs for nesting (Davidson et al., 2003; Schonberg et al., 2004); its absence from more heavily grazed forests is consistent with canopy habitat loss. Similarly, *Leptogenys* sp1 and *Platythyrea* sp1 are predators associated with humid microhabitats and dense leaf litter (Arcila-Cardona et al., 2008), resources that decline with grazing-induced litter removal and soil compaction. Incidence-based dissimilarity was driven by specialist species (*Brachymyrmex* sp1, *Acromyrmex* sp1, *Pseudomyrmex* sp1), whose plot occupancy declined along the gradient. *Pachycondyla harpax*, a predatory specialist associated with structurally intact dry forest, was not recorded in any High degradation plot, a pattern consistent with reported sensitivity to canopy disturbance in other Neotropical systems (Gove et al., 2005) and worth further investigation as a possible bioindicator of forest condition in the Tumbesian bioregion. Therefore, the monotonic decline in species richness across the grazing gradient supports our first hypothesis and falls within the range of losses reported in other Neotropical dry forests subject to canopy opening (Ríos-Casanova et al., 2006; Achury et al., 2012; Gallego-Ropero & Salguero, 2015).

However, the results contrast with those of Eldrigde et al (2020) that found that grazing favored ant species adapted to open canopy areas. In our study, three genera adapted to open habitats and considered generalist feeders (e.g., *Solenopsis* spp., *Nylanderia* spp.) increased in High degradation plots, a pattern consistent with those of Eldrigde et al (2020). Indeed, our SIMPER analyses revealed that numerical dissimilarity was driven largely by *Camponotus* spp. and *Dorymyrmex* sp1 (the former abundant throughout but variable across land degradation, the latter a thermophilic, sun-adapted *Dolichoderinae* more frequent under open, xeric conditions).

The significant decline in richness, together with the absence of significant differences in Shannon and Simpson diversity across land degradation levels, reveals an ecologically informative pattern: Low degradation retains the highest species richness but is simultaneously dominated by *Camponotus* spp. Strong numerical dominance by a single, resource-flexible species is one pattern often associated with early-stage biotic homogenization (McKinney & Lockwood, 1999; Olden et al. 2004). Although our data cannot establish whether this dominance reflects a directional homogenization process or a stable feature of structurally intact Tumbesian TDF, the dietary flexibility and resource-monopolization capacity of *Camponotus* are well documented in tropical systems (Davidson, 1998; Hölldobler & Wilson, 1990) and may simply reflect favorable conditions for a locally dominant species rather than a disturbance signature.

Similarly, grazing in our system was associated with a significant, monotonic decline in the functional space and with directional shifts in functional trait composition (smaller body size, reduced sculpturing and pilosity, and darker coloration) under High degradation. These functional shifts were supported by multiple independent lines of evidence: consistent community-weighted mean trends, a significant relationship between cattle– equine dung load and functional dispersion, and a significant ordination of trait composition along a livestock axis. These reduction in functional space with grazing are in line with previous studies (Hoenle et al. 2023), and prediction that smaller ants may be more generalist feeders (Weiser & Kaspari 2006), and that darker colorations protect ants from sunlight in open areas (Clusella-Trullas et al., 2007; Law et al., 2020).

### Caveats

Two methodological constraints should be considered when interpreting our results. First, with only three clusters per land degradation level (n = 9 clusters total), statistical power to detect compositional differences was limited, as reflected in the non-significant PERMANOVA results despite moderate-to-large effect sizes. This replication level is inherent to the logistic constraints of the study system, but it means that some real effects may have gone undetected; the significant results we do report (richness decline, CWM trait shifts) should therefore be considered conservative. Second, pitfall traps measure foraging activity rather than colony abundance, because ants are colonial central-place foragers whose trap captures depend on colony proximity and foraging range. We addressed this problem by running all analyses on both abundance and incidence data and by placing greater interpretive weight on incidence-based results where the two diverged.

### Conclusions

From a conservation perspective, these findings support the value of maintaining forests with Low degradation as refuges for taxonomically and functionally distinct ant taxa and indicate that grazing exclusion is a plausible management lever for limiting further loss of species richness and functional filtering. Because ants play key ecosystem roles in tropical dry forests, including predation of other arthropods, seed dispersal, soil aeration, and litter decomposition (Folgarait, 1998; Wirth et al., 2003), the loss of species and the directional reduction of functional trait space documented here under High degradation likely indicate a broader erosion of these processes, beyond the taxonomic decline captured by species richness alone. Therefore, forest patches with Low degradation, along with the specialist taxa they still harbor, represent not only reservoirs of biodiversity but also functional refuges for the ecological processes that underpin forest regeneration and resilience. Prioritizing the protection of these remnants and progressively excluding livestock from the most degraded stands offers a concrete and evidence-based way to safeguard both the taxonomic and functional integrity of the Tumbes dry forest, an ecosystem that has already lost more than 60% of its original extent.

## ACKNOWLEDGEMENTS

We thank the *Red Ecuatoriana de Investigación y Posgrado* (REDU) for the funding granted to the *Red de Investigación en Comunidades Ecológicas del Ecuador* (COMURED) within the framework of the 2025 Call for Research Network Registration, support that made this work possible. We extend our sincere thanks to Patricio Gusmán, Johana Gusmán, Ángel Gusmán, and Danilo Patiño for their valuable collaboration in the collection of field data. We thank two reviewers that contributed in important ways to this work.

## AUTHOR CONTRIBUTIONS

Pamela Gusmán-Montalván: Conceptualization; Investigation; Data curation; Writing – original draft; Writing – review & editing. Diego P. Vélez-Mora: Formal Analysis; Software; Visualization; Writing – original draft; Writing – review & editing. Pablo Ramón: Conceptualization; Formal Analysis. Elizabeth Gusmán-Montalván: Conceptualization; Methodology; Resources; Supervision; Writing – review & Editing. Diego Dominguez: Investigation; Data curation. David A. Donoso: Funding acquisition; Writing – review & editing.

## DATA AVAILABILITY STATEMENT

The data and R scripts supporting the results of this study are available on Figshare at https://doi.org/10.6084/m9.figshare.31768405

