## Supplemental Materials for "Goat land degradation reduces richness and reshapes the functional composition of an ant community in a tropical dry forest"

**SUPPORTING INFORMATION**

**Grazing pressure reduces ant morphospecies richness and reshapes functional trait composition in a Neotropical dry forest**

**TABLE S1.** Geographic coordinates (WGS84, decimal degrees), and grazing-pressure level of the nine sampling clusters in the Tumbesian tropical dry forest of Zapotillo, Loja Province, Ecuador. Coordinates correspond to cluster centroids.

| Cluster | Grazing-pressure level | Latitude (°) | Longitude (°) |
| --- | --- | --- | --- |
| 4 | Moderate grazing | −4.2402 | −80.3383 |
| 6 | High grazing | −4.3422 | −80.3077 |
| 7 | High grazing | −4.3422 | −80.3283 |
| 8 | Low grazing | −4.3007 | −80.3755 |
| 9 | High grazing | −4.3422 | −80.3433 |
| 10 | Low grazing | −4.3460 | −80.4025 |
| 11 | Low grazing | −4.2463 | −80.2584 |
| 15 | Moderate grazing | −4.2299 | −80.2621 |
| 16 | Moderate grazing | −4.3167 | −80.3433 |

**TABLE S2.** Post-hoc pairwise comparisons (means, Holm correction, Kenward-Roger df) of mean NDVI among the three grazing-pressure levels (linear mixed model: F₂,₆ = 8.31, p = 0.019). Mean differences are expressed as NDVI units. Estimate = mean NDVI difference between groups. P-values adjusted using the Holm method for 3 tests.

| Comparison | Estimate | SE | df | p |
| --- | --- | --- | --- | --- |
| Low vs. Moderate grazing | 0.006 | 0.0172 | 3 | 0.755 |
| Low vs. High grazing | 0.063 | 0.0172 | 3 | 0.031 |
| Moderate vs. High grazing | 0.058 | 0.0172 | 3 | 0.031 |

**TABLE S3.** Mean (± SD) Hill numbers per grazing-pressure level at the plot level (n = 9 plots per level). Differences among grazing-pressure levels were tested with linear mixed models (lmer: qD ~ Grazing_pressure_level + (1|Cluster); Kenward-Roger df; Holm correction); post-hoc comparisons with emmeans and Holm correction. q = 0: morphospecies richness; q = 1: effective number of species weighted by Shannon entropy; q = 2: effective number of species weighted by Simpson concentration.

| **Grazing-pressure level** | **q = 0 (Richness)** | **q = 1 (Shannon)** | **q = 2 (Simpson)** |
| --- | --- | --- | --- |
| Low grazing | 11.11 ± 2.21 | 3.783 ± 1.494 | 2.772 ± 1.096 |
| Moderate grazing | 10.00 ± 1.58 | 3.908 ± 0.979 | 2.909 ± 0.871 |
| High grazing | 7.22 ± 1.64 | 3.324 ± 0.849 | 2.717 ± 0.824 |

**TABLE S4.** Pairwise beta diversity decomposition between grazing-pressure levels based on presence–absence data (n = 27 plots), following Baselga (2010). β.SOR: total Sørensen dissimilarity; β.SIM: turnover component; β.SNE: nestedness component; SNE_prop: proportion of total dissimilarity attributable to nestedness (β.SNE/β.SOR).

| Pair | beta.SOR | beta.SIM | beta.SNE | SNE_prop |
| --- | --- | --- | --- | --- |
| Low vs. Moderate grazing | 0.177 | 0.099 | 0.078 | 0.41 |
| Moderate vs. High grazing | 0.227 | 0.091 | 0.136 | 0.55 |
| Low vs. High grazing | 0.277 | 0.087 | 0.190 | 0.72 |

Values computed on cluster-level presence-absence matrix (n = 9 clusters; a morphospecies scored as present if detected in ≥ 1 plot within the cluster).

**TABLE S5.** Functional trait measurements for the 22 ant morphospecies recorded in this study. For each morphospecies, at least 5 individuals were measured (45x magnification, measurement error: 0.01 mm) using an OLYMPUS SZ6 stereomicroscope. Six traits were selected for analysis: head length (HL), pilosity (PIL), sculpturing (SCUL), dominant color of head (DCH), mesosoma (DCM), and gaster (DCG).

| **Species** | **HW** | **HL** | **CL** | **ML** | **FL** | **SC** | **WL** | **PrW** | **IO** | **EL** | **BOD** | **SCUL** | **PIL** | **SPI** | **DCH** | **DCM** | **DCG** |
| --- | --- | --- | --- | --- | --- | --- | --- | --- | --- | --- | --- | --- | --- | --- | --- | --- | --- |
| *Acromyrmex* | 2.44 | 1.96 | 0.59 | 1.59 | 4.01 | 2.43 | 3.25 | 1.49 | 2.01 | 0.36 | 8 | 2 | 12 | 8 | 21 | 21 | 21 |
| *Azteca* | 1 | 1.22 | 0.31 | 0.51 | 1.22 | 0.91 | 1.2 | 0.64 | 0.62 | 0.2 | 3.56 | 3 | 39 | 0 | 16 | 17 | 15 |
| *Brachymyrmex* | 0.38 | 0.4 | 0.11 | 0.2 | 0.36 | 0.42 | 0.36 | 0.27 | 0.31 | 0.07 | 1.09 | 2 | 4 | 0 | 15 | 15 | 15 |
| *Camponotus* | 1.31 | 1.99 | 0.51 | 0.82 | 2.64 | 2.39 | 2.66 | 1.14 | 0.89 | 0.49 | 7.2 | 3 | 49 | 0 | 23 | 8 | 19 |
| *Cardiocondyla* | 0.38 | 0.51 | 0.07 | 0.22 | 0.31 | 0.31 | 0.29 | 0.24 | 0.27 | 0.11 | 1.69 | 1 | 2 | 2 | 17 | 17 | 1 |
| *Carebara* | 0.22 | 0.29 | 0.04 | 0.09 | 0.16 | 0.16 | 0.29 | 0.16 | 0 | 0 | 1.07 | 2 | 10 | 0 | 16 | 16 | 16 |
| *Cephalotes* | 1.24 | 0.82 | 0.02 | 0.34 | 0.6 | 0.4 | 0.94 | 0.8 | 1.02 | 0.36 | 3.3 | 2 | 0 | 14 | 2 | 2 | 2 |
| *Crematogaster* | 0.86 | 0.87 | 0.24 | 0.37 | 0.77 | 0.58 | 0.81 | 0.46 | 0.67 | 0.18 | 3.23 | 1 | 2 | 2 | 24 | 23 | 12 |
| *Cyphomyrmex* | 0.62 | 0.67 | 0.22 | 0.38 | 0.73 | 0.47 | 0.87 | 0.47 | 0.51 | 0.13 | 2.4 | 3 | 0 | 8 | 8 | 8 | 8 |
| *Labidus* | 0.93 | 1.04 | 0.09 | 0.58 | 1.2 | 0.73 | 1.4 | 0.62 | 0.51 | 0.04 | 3.96 | 2 | 42 | 0 | 22 | 21 | 17 |
| *Dorymyrmex* | 0.71 | 0.96 | 0.18 | 0.42 | 0.98 | 0.87 | 1.02 | 0.49 | 0.36 | 0.22 | 2.69 | 1 | 0 | 1 | 15 | 15 | 1 |
| *Leptogenys* | 0.58 | 0.87 | 0.18 | 0.4 | 0.89 | 0.76 | 1.33 | 0.49 | 0.44 | 0.11 | 4.1 | 1 | 39 | 0 | 22 | 22 | 22 |
| *Neivamyrmex* | 0.47 | 0.44 | 0.07 | 0.31 | 0.33 | 0.36 | 0.49 | 0.33 | 0.42 | 0.04 | 1.6 | 3 | 19 | 2 | 24 | 24 | 24 |
| *Nesomyrmex* | 0.44 | 0.53 | 0.25 | 0.8 | 0.44 | 0.25 | 2.1 | 0.75 | 0.89 | 0.29 | 1.2 | 2 | 17 | 3 | 21 | 22 | 24 |
| *Nylanderia* | 0.49 | 0.63 | 0.17 | 0.32 | 0.66 | 0.74 | 0.72 | 0.39 | 0.3 | 0.16 | 2.03 | 1 | 6 | 0 | 8 | 8 | 8 |
| *Odontomachus* | 2.01 | 3.93 | 0.25 | 1.52 | 2.77 | 2.42 | 3.09 | 1.13 | 1.43 | 0.44 | 9.1 | 2 | 73 | 1 | 1 | 16 | 1 |
| *Pachycondyla* | 1.71 | 2.48 | 0.4 | 1.16 | 1.76 | 1.42 | 2.77 | 1.27 | 1.38 | 0.31 | 10 | 2 | 44 | 0 | 1 | 1 | 1 |
| *Pheidole* | 0.73 | 0.67 | 0.24 | 0.47 | 0.73 | 0.67 | 0.93 | 0.42 | 0.64 | 0.13 | 2.86 | 1 | 21 | 2 | 24 | 24 | 24 |
| *Platythyrea* | 1.09 | 0.82 | 0.4 | 0 | 0 | 0 | 0 | 0 | 0 | 0 | 0 | 3 | 42 | 3 | 8 | 24 | 24 |
| *Pseudomyrmex* | 1.09 | 0.85 | 0.2 | 0.76 | 1.11 | 1.02 | 1.62 | 0.62 | 0.78 | 0.16 | 4.87 | 2 | 36 | 4 | 1 | 1 | 24 |
| *Rogeria* | 0.47 | 0.69 | 0.11 | 0.29 | 0.36 | 0.38 | 0.6 | 0.36 | 0.42 | 0.07 | 1.91 | 1 | 21 | 2 | 17 | 17 | 15 |
| *Solenopsis* | 0.29 | 0.38 | 0.07 | 0.18 | 0.24 | 0.22 | 0.38 | 0.2 | 0.26 | 0.03 | 1.33 | 1 | 8 | 0 | 18 | 18 | 18 |

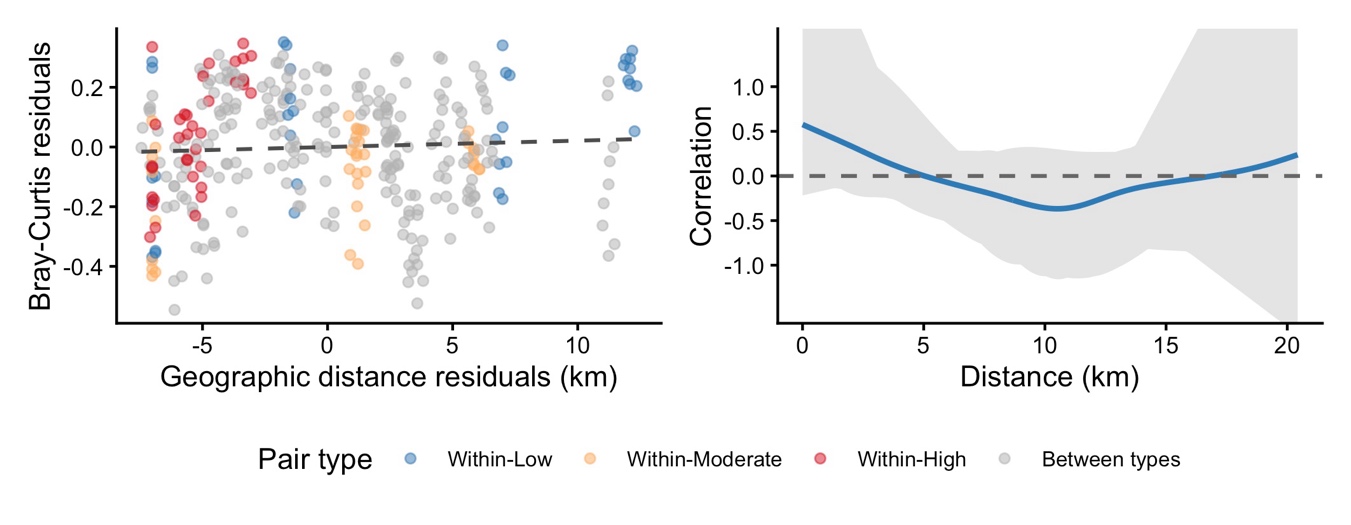

**FIGURE S1**. Spatial independence of ant community composition across the grazing gradient. (a): Partial Mantel test scatter plot showing Bray-Curtis compositional dissimilarity residuals versus geographic distance residuals after controlling for grazing-pressure level (r = 0.055, p = 0.238); each point represents one of the 351 pairwise plot comparisons, coloured by pair type (Within-Low grazing, Within-Moderate grazing, Within-High grazing, Between types); the dashed line shows the OLS regression through the residuals. (b) Spline correlogram of the first residual axis of a dbRDA (Bray-Curtis; standardised scores) as a function of geographic distance between plot centroids (km). The blue line shows the observed spatial correlation; the grey band represents the 95% pointwise confidence interval derived from 999 bootstrap permutations (width reflects reduced plot-pair availability at extreme distances). Positive autocorrelation at distances < 0.5 km corresponds to within-cluster plot spacing (310–438 m centroid-to-centroid) and reflects the nested sampling design rather than treatment confounding. At all inter-cluster distances (> 0.5 km), the confidence band includes zero, confirming spatial independence at the scale of treatment assignment.

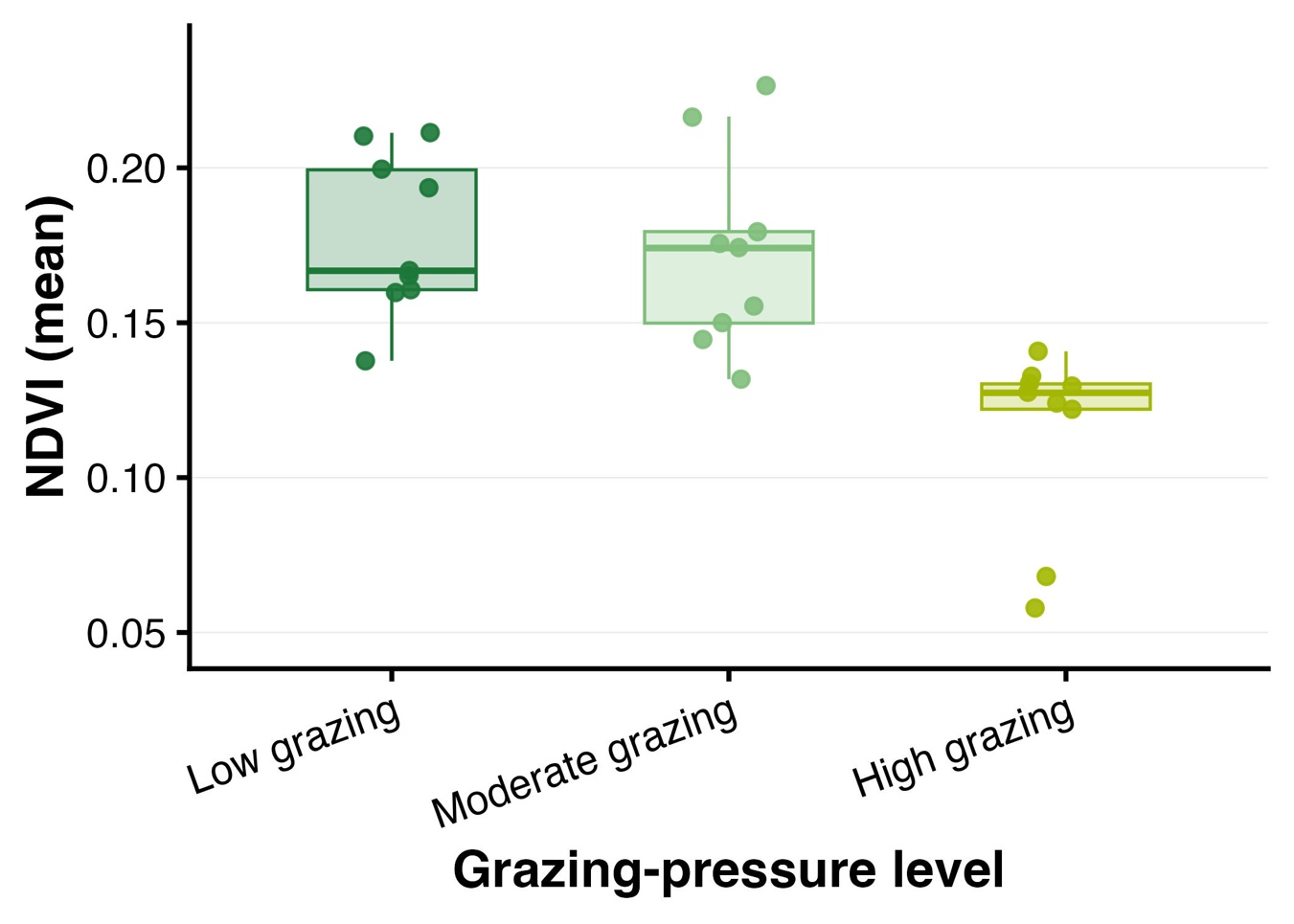

**FIGURE S2.** Mean NDVI per grazing-pressure level extracted from Sentinel-2 imagery (November 2018, COPERNICUS/S2_HARMONIZED, Level 1C). Boxplots show median and interquartile range; points show individual plot values (n = 9 per level).

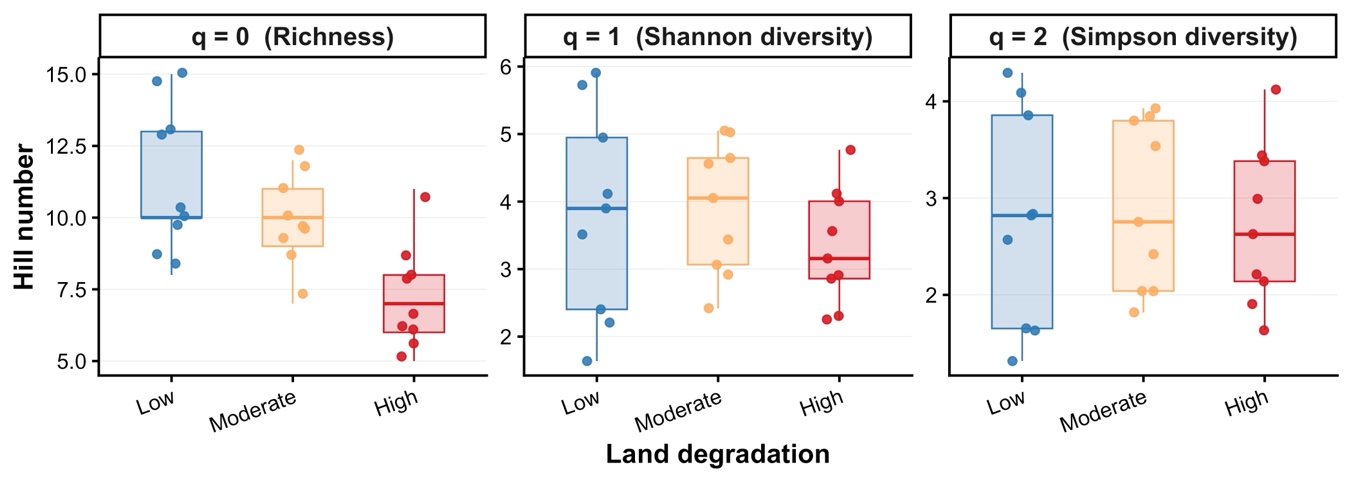

**FIGURE S3.** Hill numbers for ant morphospecies richness and diversity at the cluster level (n = 9 clusters, 3 per grazing-pressure level). q = 0: morphospecies richness; q = 1: Shannon diversity; q = 2: Simpson diversity. Only q = 0 differed among grazing-pressure levels (linear mixed model: F2,6 = 5.64, p = 0.042). White diamonds indicate cluster-level means (n = 3 per grazing-pressure level), representing the unit of treatment replication.

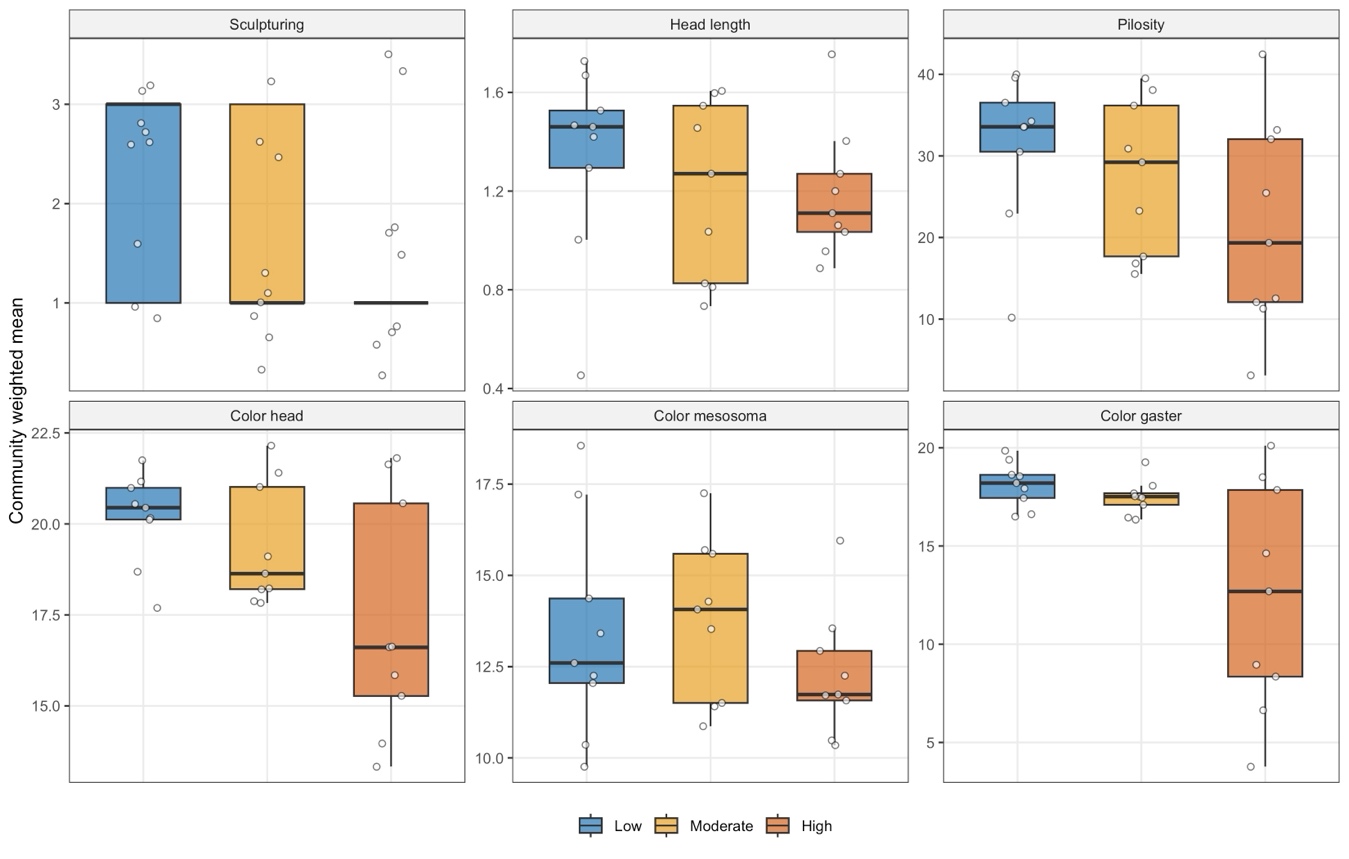

**FIGURE S4.** Community-weighted means (CWM) of six functional traits across the grazing gradient: head length (HL), pilosity (PIL), sculpturing (SCUL), and dominant color of head (DCH), mesosoma (DCM), and gaster (DCG). Boxplots show median and interquartile range; points show individual plot values (n = 9 per level).

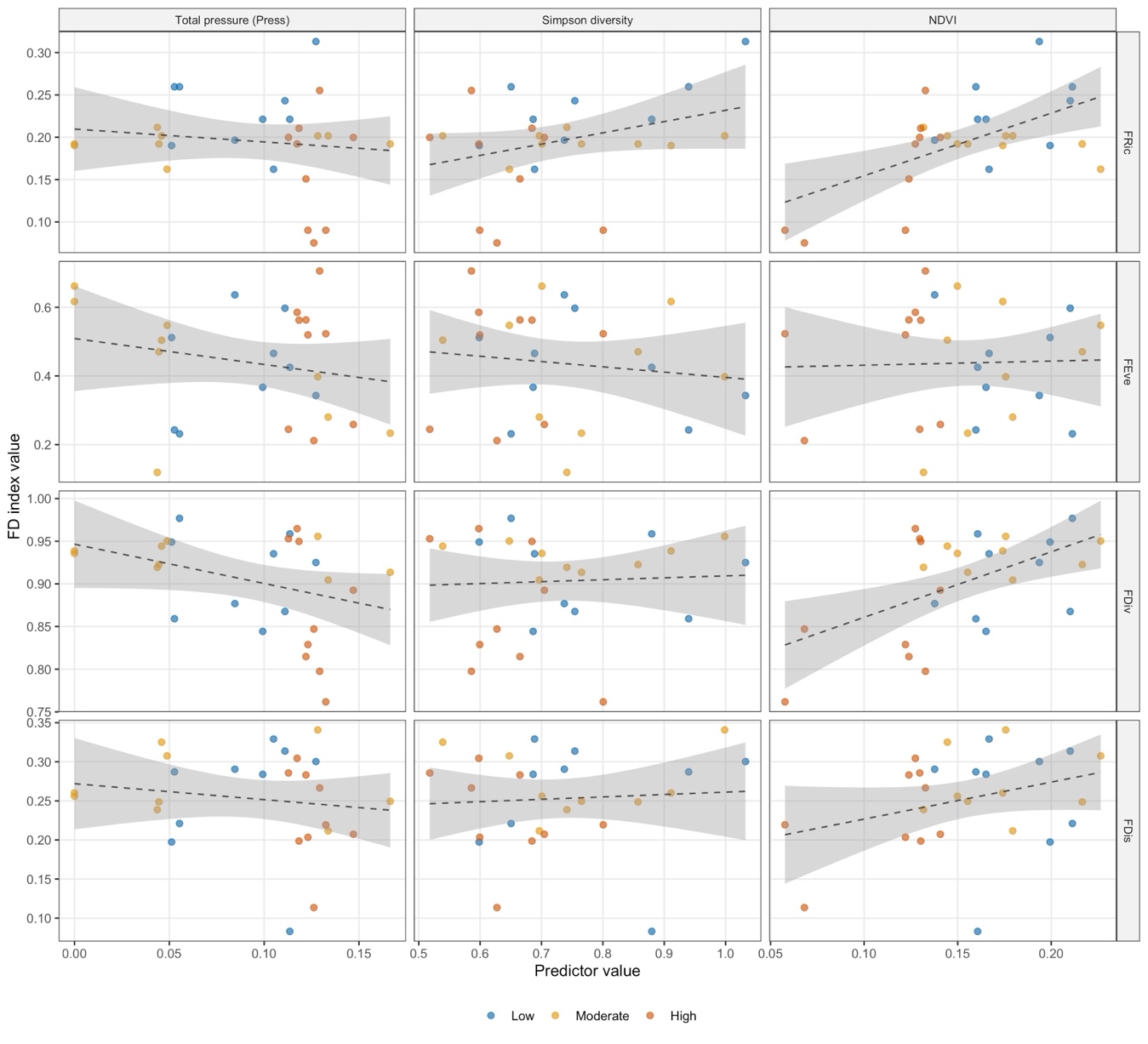

**FIGURE S5.** Functional diversity indices (Functional Richness, Functional Evenness, Functional Divergence, Functional Dispersion) plotted against three continuous predictors: total livestock pressure (Press), tree species diversity (Simpson), and normalized difference vegetation index (NDVI). Each point represents one plot (n = 27); colors indicate grazing-pressure level. Linear trend lines are fitted per predictor.

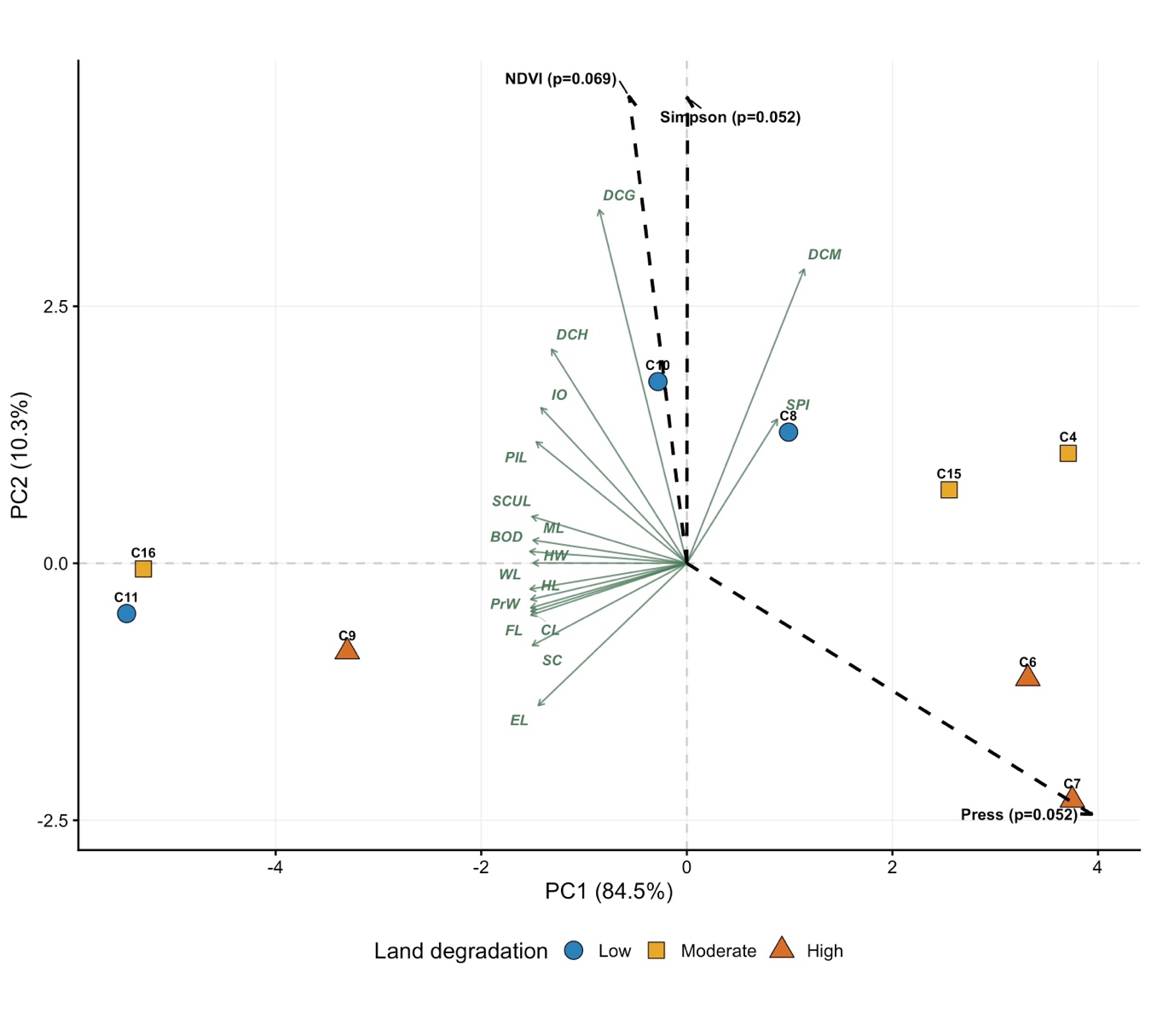

**FIGURE S6.** Principal component analysis (PCA) biplot of cluster-level CWM of all 17 functional traits (n = 9 clusters). This supplementary PCA confirms that the six-trait subset used in the main text captures the dominant functional gradients (main-text PCA: PC1 = 76.6%, PC2 = 21.8%; this PCA: PC1 = 84.5%). Environmental predictors (Press, Simpson, NDVI) projected via envfit (999 permutations).

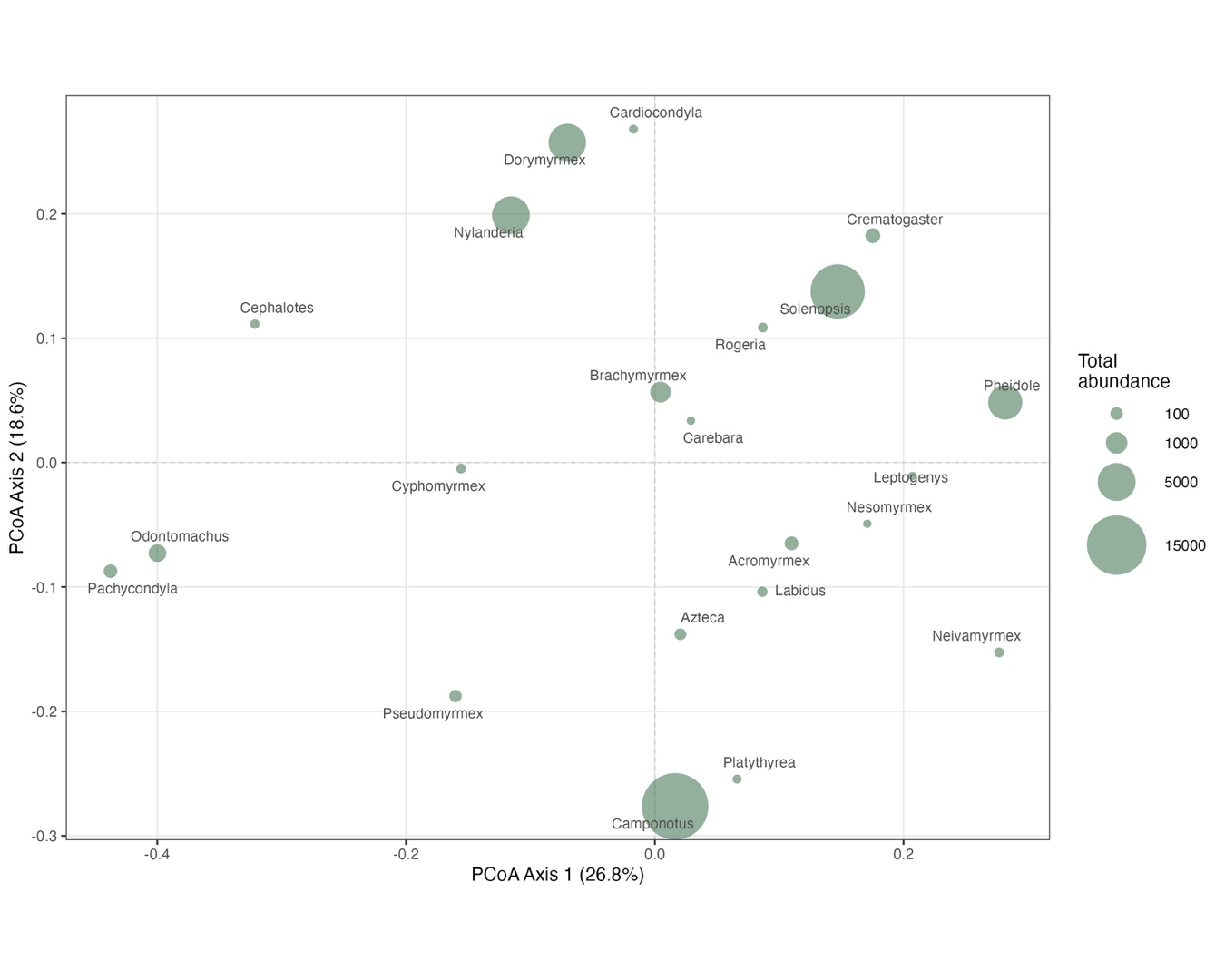

**FIGURE S7.** Species-level functional space based on PCoA of Gower distances among 22 genera using six functional traits. Bubble sizes are proportional to morphospecies abundance within each level. The occupied functional space contracts progressively from low grazing (all 22 morphospecie) to high grazing (dominated by four generalist genera).
